# Impaired humidity sensing reduces tick survival by preventing water homeostasis

**DOI:** 10.1101/2024.11.06.618166

**Authors:** Melissa R. Uhran, Kosisochukwu Onyeagba, Shania M. Sanderson, Syeda Farjana Hoque, Melissa Kelley, Evan S. Smith, Kennan Oyen, David A. Lewis, Ayesha Benton-Anderson, Thomas Arya, Arturo Ledezma, Haidar Almatar, Jasmine Kennedy, Ronja Frigard, Christopher J. Holmes, Shyh-Chi Chen, Pia U. Olafson, Holly D. Gaff, Joshua B. Benoit

**Author notes:** Author for correspondence: Joshua B. Benoit, Department of Biological Sciences, University of Cincinnati, Cincinnati, OH.

## Abstract

Off-host periods are critical for ticks, representing a time when environmental stress, particularly dehydration, can impact tick survival. To prevent dehydration, ticks must be able to detect and move into high humidity areas to allow for water vapor uptake. Ionotropic receptor 93a (Ir93a), which is highly expressed in the front forelegs (location of Haller’s organ), increases expression following dehydration, suggesting a role for the forelegs in humidity detection. Although the Haller’s organ is suggested as the site of humidity detection in ticks, humidity detection has not been extensively examined for ticks. Here, we assessed the ability of ticks to sense humidity and how altered humidity detection impacts tick survival by manipulating the Haller’s organ. Permanent (cutting or heat ablation) or temporary blocking (silicone covering) of the Haller’s organ impairs the ability of the American dog tick, *Dermacentor variabilis,* to rest in areas necessary to maintain hydration. Impaired detection of humidity did not impact tick survival when individuals were held under stable optimal conditions. Still, variable conditions (low and high humidity gradient) reduced survival through dehydration stress and decreased energy reserves, as chronic water vapor uptake is energetically expensive. Field validation of these studies in *D. variabilis* confirmed that humidity detection is critical to tick survival. Lastly, modeling indicates impaired humidity detection will reduce questing tick populations, specifically in the adult stages. These studies confirm that the Haller’s organ is critical for humidity sensing in ticks. Without this ability, tick survival will be impaired by potential dehydration or more rapid depletion of energy reserves.

## Introduction

Ticks act as vectors for a myriad of disease causing pathogens but spend the majority of their lives between blood meals, especially if the tick is a three host species. During off-host periods, ticks are at risk for dehydration due to their small body size and consequent high surface-area-to-volume ratio (Benoit and Denlinger, 2010), and maintaining water balance is critical for tick survival (Needham and Teel, 1991; Rosendale et al., 2017). Ticks prevent dehydration through adaptations to conserve water or rely on the acquisition of exogenous water (Benoit and Denlinger, 2010; Needham and Teel, 1991). Water is replenished by an active, solute-driven process of water vapor absorption (Gaede and Knülle, 1987). This process of obtaining water from the air reduces lipid reserves but prevents premature death from dehydration (Randolph and Storey, 1999; Rosendale et al., 2017). As ticks feed only on blood, nutrient reserves are limited to those obtained from the egg and subsequent blood meals (Rosendale et al., 2019). Although ticks show a remarkable ability to withstand prolonged starvation, partly through low metabolic rates (Lighton and Fielden, 1995), exhaustion of energy reserves is a significant cause of mortality under field conditions (Nieto et al., 2010). Thus, understanding specific factors that allow tick survival between blood meals is critical to potential pathogen transmission.

The ability of ticks to survive relies on locating humidities above critical equilibrium humidity (CEH) (Gaede and Knülle, 1987; Maldonado-Ruiz et al., 2020; Yoder et al., 2012), which requires the ability to sense humidity. The primary sensory organ of ticks is the Haller’s organ, a unique sensory structure situated exclusively on the 1st pair of legs (Carr et al., 2017). This organ is expected to act as the functional equivalent of insect antennae in relation to the sensation of the environment but is morphologically dissimilar to the antennae. In general, little is known about sensory perception in ticks. Recent studies have confirmed that the sensory structures of the Haller’s organ responded directly to specific host cues such as humidity, looming stimulus, and olfactory cues (Josek et al., 2021). Transcriptome analyses of this organ in the black-legged tick, *Ixodes scapularis*, revealed that specific ionotropic receptors are highly expressed in the foreleg compared to other legs (Josek et al., 2018). Ionotropic receptors have been extensively studied in numerous insect systems and act as ligand-gated channels for chemosensation (Enjin et al., 2016; Graeve et al., 2022; Laursen et al., 2023). Specifically, there is an enrichment for Ir93a and Ir25a in the forelegs, which are differentiated from the other legs by the presence of the Haller’s organ. Recent studies on insect systems have established that Ir93a is a key factor in humidity detection for arthropod systems (Enjin et al., 2016; Graeve et al., 2022; Laursen et al., 2023).

In this study, we examined humidity detection in American dog ticks, specifically the role of the Haller’s organ in this process, and how this detection impacts the survival of ticks under variable environments. Ticks preferred to locate and reside in more humid microenvironments, but this was impaired by interference with the Haller’s organ. Transcript expression patterns of Ir93a show that this receptor is expressed highly in the forelegs of ticks, which is the site of the Haller’s organ. Haller’s organ interference resulted in increased mortality, predominantly through the rapid decline in lipid reserves and increased starvation-induced death when exposed to dry conditions. Field observations support that interference with the Haller’s organ will prevent tick survival, highlighting the importance of this process. These studies confirm that the Haller’s organ, likely through a contribution from Ir93a, is critical in humidity detection. This humidity detection is crucial for tick survival under dehydrating and variable conditions by preventing starvation-induced death. Modeling of impaired humidity suggests that tick populations would decline if periods of drought were to occur. Altogether, we demonstrate the importance of the Haller’s organ in humidity detection of ticks and the critical role of humidity detection for tick survival.

## Materials and Methods

### Ticks

Adult *D. variabilis* were acquired from Ectoservices (Cary, NC), and adult female *Amblyomma americanum* were acquired from the USDA Knipling-Bushland U.S. Livestock Insects Research Laboratory. Ticks from Ectoservices were fed on sheep (*Ovis aries*), and those from the USDA were fed on cattle (*Bos taurus*). Before experimentation, ticks were stored at 24-25°C and 12:12 L:D light cycle. Relative humidities were held at 85% or 93% relative humidity (RH) with saturated salt solutions (Winston and Bates, 1960). All ticks were two weeks of age when used in the studies and only female ticks were used, with the exception of the transcript assessment. The majority of the studies were conducted on *D. variabilis* with some select studies performed on *A. americanum* (survival analyses).

### Interference with Haller’s organ

To prevent Haller’s organ function, we performed a series of manipulations that consisted of wax covering, heat ablation, and leg removal (Fig. 1). Sets of legs are numbered anterior to posterior for this paper. In leg removal, either the first or second set of legs were removed; this allows for comparison of ticks with and without their Haller’s organ. In heat ablation, a heated metal probe was used for the cauterization of the Haller’s organ or a similar location on the second set of legs. This allowed for the interference of the Haller’s organ with the removal of the underlying nervous tissue. In wax covering, the tick forelegs were covered with warm dental silicon (Defend). These wax coverings can later be removed to allow humidity detection recovery. Following interference with the legs and Haller’s organs, ticks were held at 100% RH (two days) to allow for recovery before use in the experiments within 3-5 days.

**Figure 1.**
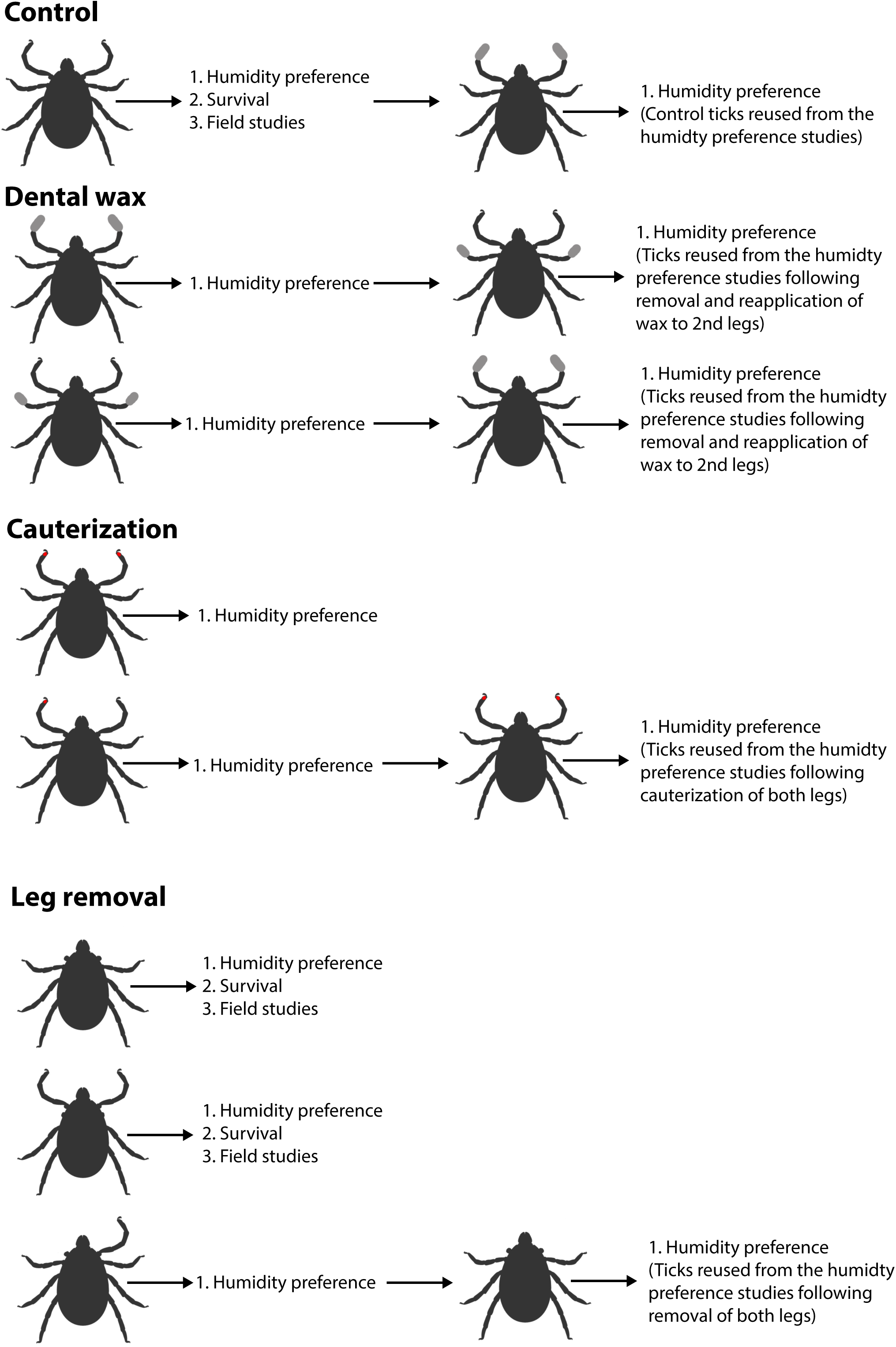
Schematics for leg interference. Schematics of the processes that impact Haller’s organ functionality include temporary impairment by blocking with dental wax that can be removed, interference through cauterization, and permanent removal through amputation of the legs.

### Humidity preference assays

To determine the ability to detect and subsequently start quiescent periods at different humidities, we developed a two-choice assay. A 42 cm polycarbonate tube (2.54 cm i.d.) was capped with two *Drosophila* vials (25 x 95 mm) that were filled with calcium carbonate (Drierite, Xenia, OH) to generate low relative humidity (0-10% RH), and saturated sodium chloride (75% RH) to generate medium relative humidity, or sponges soaked with deionized water (95-100% RH) to generate high relative humidity. Ticks were added into the 42 cm tube in the middle and each side was capped with different vials containing the two solutions, yielding an attraction distance of 40 cm. Tick locations were assessed at 24 hours as this would be the location most likely for the tick to rest for more extended periods. Ticks that did not survive 24 hours were removed from these analyses. Values were converted to a humidity attraction value based on the distance from the dry side, thus, if a tick was 40 cm from the dry side, it would convert to a humidity attraction value of 1.

### Lipid content

Lipid reserve levels were monitored during the course of laboratory-based survival studies under both constant and variable relative humidities. Following removal for experiments, ticks were stored at −70°C until samples were used in assays. Pairs of ticks were homogenized in 500 μl of STE buffer (250 mM NaCl, 10 mM Tris, 5 mM EDTA, pH 8.3) with 2% Na_2_SO_4_ using a BeadBlaster 24 microtube homogenizer (Benchmark Scientific, Edison, NJ, USA). Total lipid content was determined based on methods from van Handel (Van Handel, 1985) that have been modified for analysis of ticks (Rosendale et al., 2017; Rosendale et al., 2019). In brief, the homogenate (150 μl) was combined with 250 μl 1:1 chloroform:methanol, vortexed, centrifuged to remove debris, and the solvent was evaporated at ∼90 °C. The lipid-enriched residue was dissolved with concentrated H_2_SO_4_ by heating for 30 minutes at ∼90 °C. Vanillin-phosphoric acid reagent was added and absorbance was measured at 525 nm after 10 minutes. Lipid levels were compared to a standard of canola oil dissolved in chloroform.

### Water content determination

Standard methods for determining the water content of the ticks were used and based directly on those previously reported in ticks and other arthropod systems (Bailey et al., 2023; Benoit et al., 2007; Yoder et al., 2012). Ticks were weighed using an electrobalance (SD ± 0.2 μg precision and ± 6 μg accuracy at 1 mg based on five separate mass measurements using a 1-mg weight at the 200-mg range; CAHN; Ventron Co., Cerritos, CA, U.S.A.). The fresh mass was determined immediately after removal from the experiment. The dry mass was determined after drying each tick at 0% RH and 70 °C (drying oven) for seven days. The tick was reweighed for at least three days to ensure the dry mass remained constant. Water mass is the difference between the fresh and dry mass and the proportion of water content is the water mass/fresh mass.

### Laboratory survival assay

Survival assays were conducted on ticks in variable and constant relative humidities. Ticks were held at either 85% RH for constant humidity exposure and monitored for survival until 100% mortality occurred. Survival was assessed by observing tick movement following a pulse of breath applied by the experimenter. Tick survival under variable humidities was accomplished using the same setup as in the humidity preference assays with a low and a high RH that the ticks could access based on their choice. Vials to generate humidities were changed every two days. All experiments were conducted at 24-25°C and 12:12 L:D. A subset of ticks was collected at specific weekly intervals for the assessment of water balance metrics as proxies for hydration levels and lipid reserves as an assessment of nutrient reserve utilization as discussed previously.

### Field survival assays

Along with laboratory-based survival studies, we conducted field survival assessments at the University of Cincinnati Center for Field Studies (Harrison, OH). Groups of ten ticks were housed in small cages (Bioquip, 10 × 10 × 10 cm) and placed randomly under tree cover to prevent direct sunlight onto the cages. Five cages were evaluated for each treatment from April - November 2021. The treatments examined were the front legs removed, 2nd legs removed, and control (no treatment). Tick survival was evaluated each week until mortality was 100% or the first freezing event occurred in late November. At select points (two, ten, and twenty weeks), ticks were removed to assess hydration levels and lipid reserves.

### Quantitative PCR of American dog tick forelegs

To assess if enrichment is localized in the forelegs when compared to the body and other legs, we used qPCR to determine relative transcript levels for Ir93a. qPCR analyses were conducted using previously developed methods (Rosendale et al., 2016). Additionally, ticks were collected before or after dehydration (20% water loss after five days), and after rehydration following a two day period at 97% RH (Rosendale et al., 2016; Rosendale et al., 2017). Males and females at one month of age were used. RNA was extracted as described previously for three independent biological replicates. Complementary DNA (cDNA) was generated with a DyNAmo cDNA Synthesis Kit (Thermo Scientific). Each reaction used 250 ng RNA, 50 ng oligo (dT) primers, reaction buffer containing dNTPs and 5 mmol•l^−1^ MgCl_2_, and M-MuLV RNase H+ reverse transcriptase. KiCqStart SYBR Green qPCR ReadyMix (Sigma Aldrich, St Louis, MO, USA) along with 300 nmol l^−1^ forward and reverse primers, cDNA diluted 1:20, and nuclease-free water were used for all reactions. Primers were designed using Primer3 based on contigs obtained from a previous transcriptome analysis (Rosendale et al., 2022), F-ATGACGAACAGCCACCAGAA and R-GCTTCGGGGCAATCATTCAA. qPCR reactions were conducted using an Illumina Eco quantitative PCR system. Five biological replicates were examined for each Sample. Expression levels were normalized using the ΔΔCq method as previously described (Finch et al., 2020; Rosendale et al., 2016; Rosendale et al., 2019; Rosendale et al., 2022).

### Modeling of tick survival following impaired humidity detection

To assess the potential long-term impact of reduced survival, we used a life-history matrix model to estimate the predicted reduction in tick populations (Sanderson et al., 2024). This American dog tick model (ADTSIM 2.0) was based on a previous model for *D. variabilis* populations (Mount and Haile, 1989) and was updated with more recent research similar to the recently updated LYMESIM model for *Ixodes scapularis* (Gaff et al., 2020). Similar to the updates for LYMESIM, ADTSIM was updated to include a finite lifespan and a survival penalty for questing. We used the 2007-2016 average weather and climatic conditions from Norfolk, Virginia, USA for baseline metrics. Three model scenarios were used with changes to the survival rates of questing ticks: The first had the standard survival rates for all life stages of ticks, the second had reduced survival for just the adult life stage, and the third had reduced survival in all life stages.

### Statistics and replication

R was utilized for all statistical analyses (Core Team and Others 2013). All data were examined for normal distribution before analyses. ANOVA was used to compare treatments and specific differences between each were established with Tukey’s post hoc test. Generalized additive models (GAMs) were generated and smoothed with the MGCV package (Wood, 2004; Wood, 2006; Wood, 2017) in R to analyze survival data. GAMs were compared pairwise with Tukey’s honestly significant difference post-hoc tests through the emtrends function of the emmeans package (Lenth, 2020).

## Results

### Expression patterns suggest Ir93a is in the forelegs

Previous studies have confirmed that a specific ionotropic receptor (Ir93a) is the critical component of humidity detection and is present in ticks (Laursen et al., 2023). Targeted expression analyses of *Ir93a* within specific legs confirmed that the expression levels are highly enriched in the forelegs and are enriched in foreleg compared to the remainder of the tick body (Fig. 2). Lastly, dehydration yielded a significant increase in Ir93a, suggesting a responsiveness to prolonged exposure to dry conditions (Fig. 2). These experiments provide strong evidence that Ir93a is likely involved in humidity detection in the Haller’s organ.

**Figure 2.**
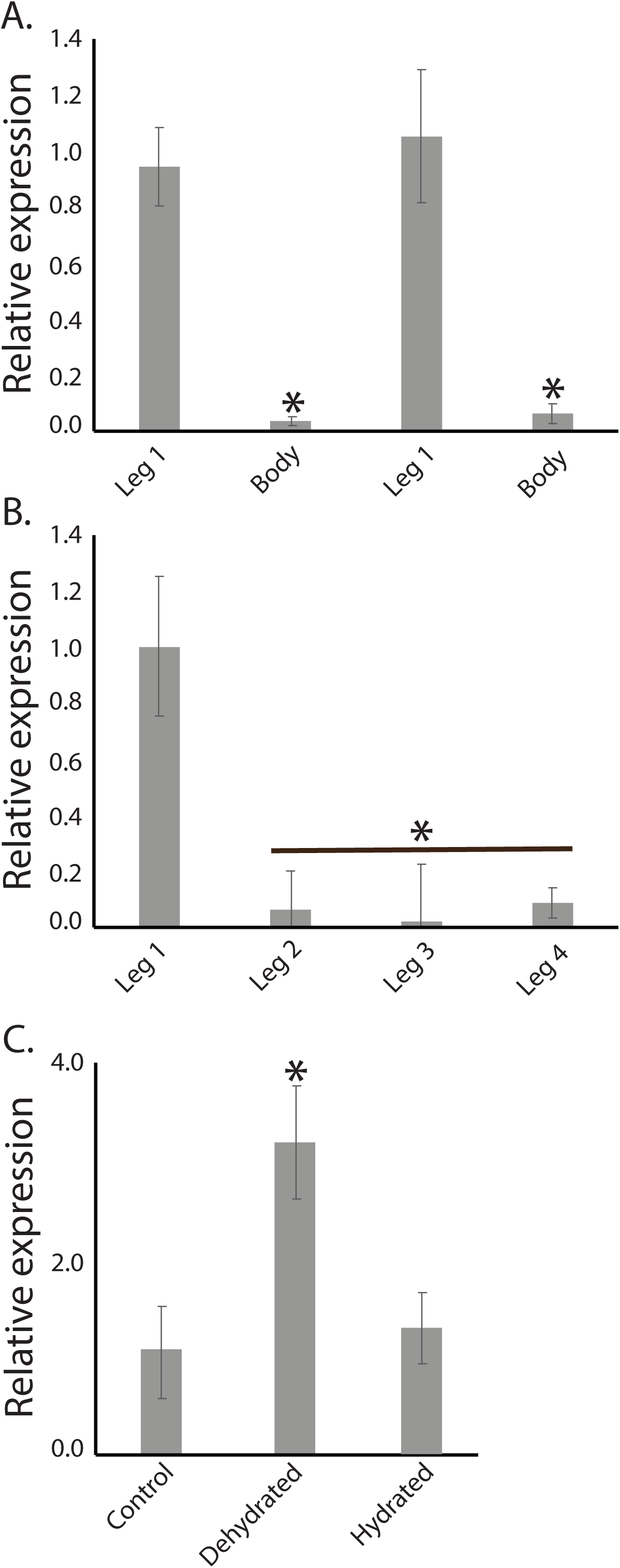
Expression analysis of ionotropic receptor 93a in the American dog tick. A. quantitative PCR of forelegs compared to the body of males (left) and females (right). N = 5 for each sample. *, significantly different, t-test, P < 0.05. B. quantitative PCR of each leg set. N = 5 for each leg set, *, significantly different, ANOVA, P < 0.05. C. Expression patterns before and after dehydration and hydration. N = 5 for each leg set, *, significantly different, P < 0.05, Tukey’s ANOVA.

### Haller’s organ interference impacts humidity detection

When American dog ticks were allowed access to low or high humidity, individuals with intact forelegs predominantly rested at higher relative humidities (Fig 3). Removal of the first set of legs had a significant impact on humidity detection, in that there was no preference to rest at a low or high humidity (Fig. 3). Removal of a single foreleg allowed the ticks to choose and remain at the higher humidity. A similar result was observed when Haller’s organ was cauterized on one leg (Fig 3). Lastly, covering the tip of the front legs yielded ticks that did not prefer high humidity, which could be recovered after wax removal. These studies suggest that the forelegs, most likely the Haller’s organ, are involved in tick humidity detection and preference.

**Figure 3.**
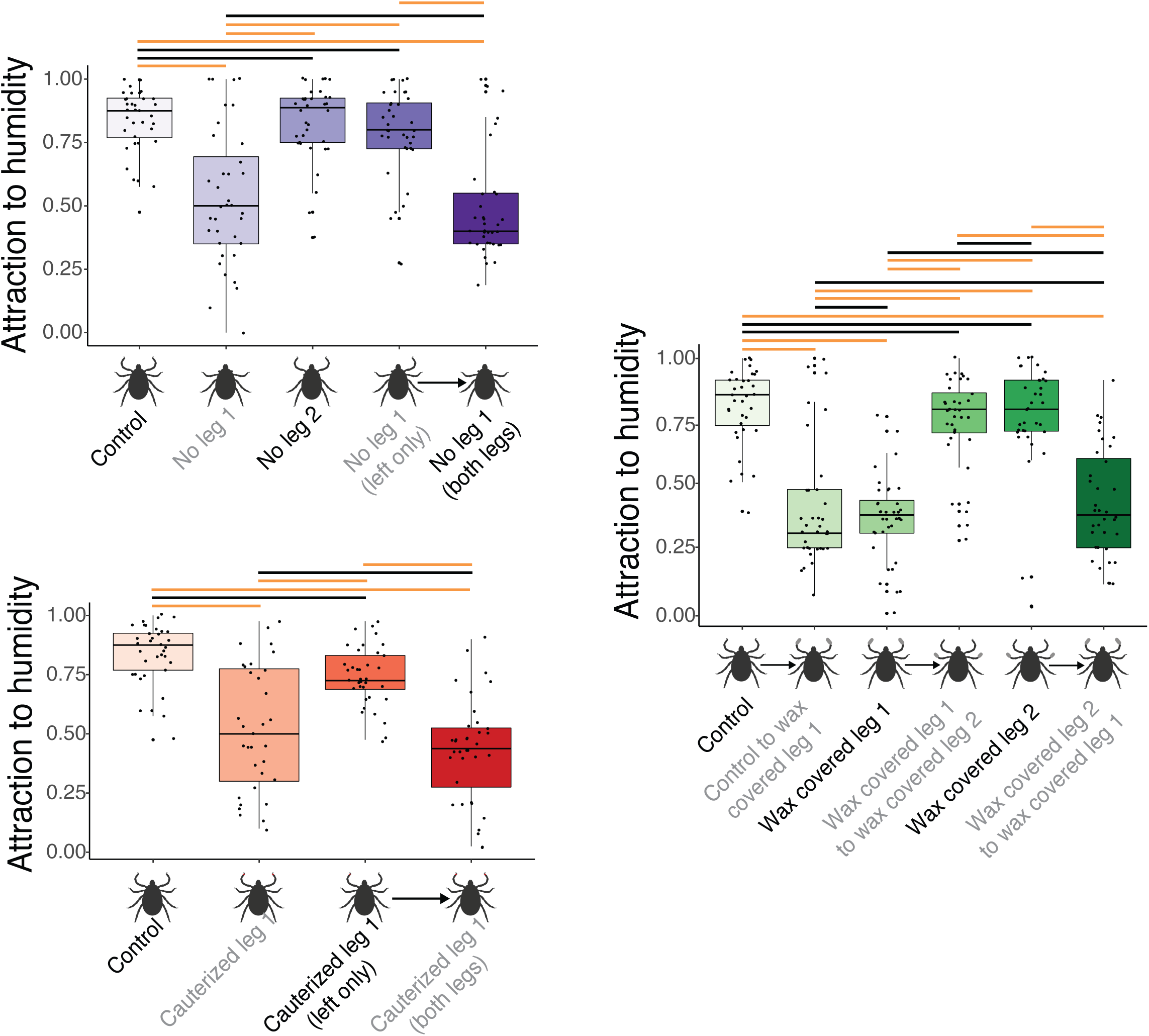
Humidity detection in adult female *Dermacentor variabilis* with and without impaired humidity detection. Top left (Purple). Leg treatment occurred three days before humidity attraction assay. Bottom left (Red). Cauterization of Haller’s organ on the forelegs three days before attraction assay. Right (Orange). Wax covering of the forelegs. The orange bars above indicate significant differences between the treatments (ANOVA, Tukey’s, P < 0.05). Black bars indicate that there are no significant differences between treatments.

### Reduced humidity detection impacts survival under variable humidities

As the removal of the legs was a functional mechanism to reduce tick humidity detection, we used this treatment to evaluate how this impacts prolonged survival. When the ticks were held at stable conditions (85% RH), there were only minimal differences in survival between ticks without the first or second legs (Fig. 4). When allowed a choice between low and high humidity, there was a significant reduction in survival for ticks lacking front legs compared to those with their full set of legs or those with the second set of legs removed (Fig. 4).

**Figure 4.**
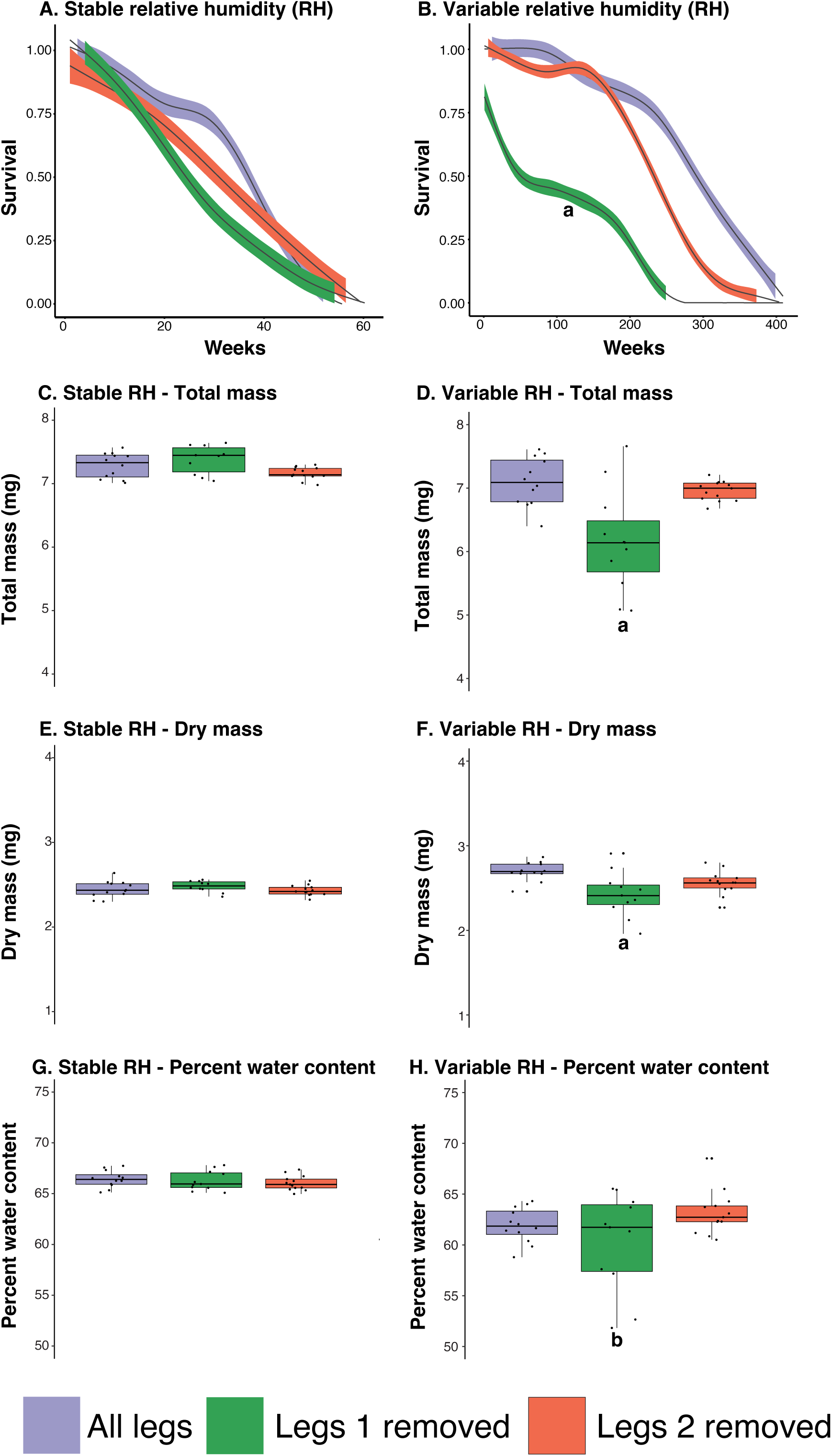
Survival, mass change, and water content of American dog ticks after foreleg removal under constant or variable relative humidities. Stable relative humidity was achieved with a saturated salt solution (potassium nitrate) to generate 93% RH. Variable humidity was generated by placing the ticks in the humidity attraction tube, where one end is ∼20% RH and the other is 95% RH. A. Survival under constant RH (N = 8 groups of 10 ticks). B. Survival under variable RH (N = 8 groups of 10 ticks). The generalized additive model for survival shows a significant between the removal of the first legs compared to the control and second leg removal (P < 0.001). C. Total mass under constant RH (N = 8 groups of 10 ticks). D. Total mass under variable RH (N = 8 groups of 10 ticks). E. Dry mass under constant RH (N = 8 groups of 10 ticks). F. Dry mass under variable RH (N = 8 groups of 10 ticks).). G. Percent water content under constant RH (N = 8 groups of 10 ticks). H. Percent water content under variable RH (N = 8 groups of 10 ticks). All mass and water content changes were determined after 12 weeks under constant or variable conditions. A, indicates a significant difference between ticks with all legs, forelegs missing, and second legs missing. B, indicates a substantial increase in variation between groups (P < 0.05, Tukey’s ANOVA).

When mass changes were assessed, there was a substantial decline in the total and dry mass following the removal of the first set of legs compared to the controls (Fig. 4C-H). Of interest, there was a significant increase in variation in the total mass and water mass when humidity sensing legs were removed, suggesting an increase in variation of the water content. Lipids levels were more rapidly utilized under variable conditions compared to stable humidity conditions (Fig. 5). Similar trends in survival were obtained for the lone star tick, *A. americanum* (Fig. S1, S2). These studies suggest that removing the legs with the Haller’s organ will impact tick survival under variable humidity conditions.

**Figure 5.**
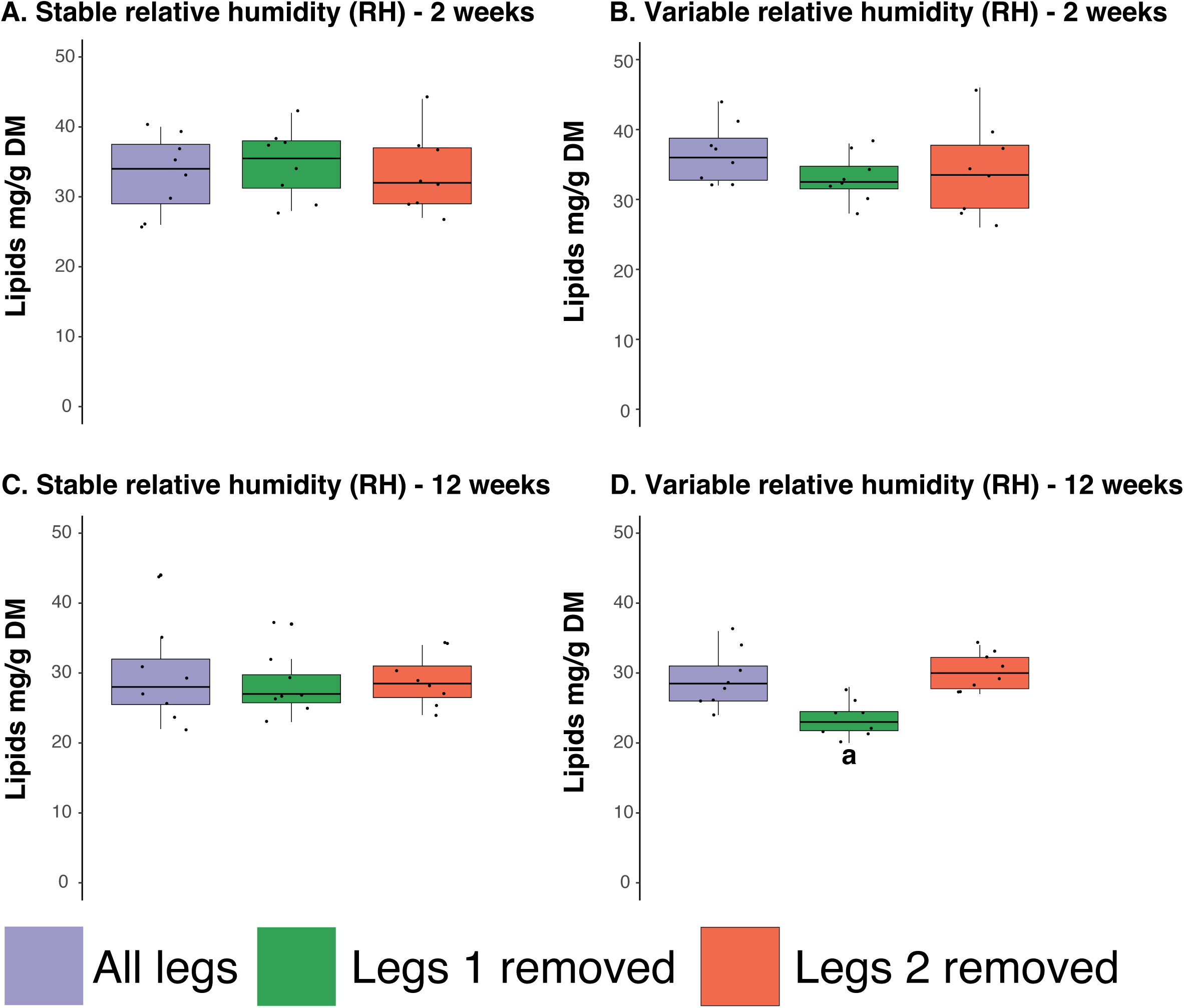
Lipid content under constant or variable relative humidities. Stable relative humidity was achieved with a saturated salt solution (potassium nitrate) to generate 93% RH. Variable humidity was generated by placing the ticks in the humidity attraction tube, where one end is ∼20% RH and the other is 95% RH. A-B. After two weeks of treatment, C-D, lipid content under stable and variable RHs. Lipid content under stable and variable RHs after twelve weeks of treatment, a, indicates a significant difference between ticks with all legs, forelegs missing, and second legs missing (P < 0.05, Tukey’s ANOVA). Sample sizes, N = 8-10.

### Tick survival is drastically reduced under field conditions due to increased metabolism

As variable conditions were shown to impair survival, we assessed the survival of ticks under field conditions. Similar to the laboratory studies, we show that the removal of the forelegs leads to a reduction in survival (Fig. 6) compared to ticks with all legs intact or with their second set removed. Measurement of lipid reserve levels after 2, 10, and 24 weeks under field conditions indicated the lipid reserves decline more rapidly in ticks without their forelegs (Fig. 5). These studies indicate that impaired humidity detection will lead to reduced survival over extended periods due to more rapid utilization of lipid reserves.

**Figure 6.**
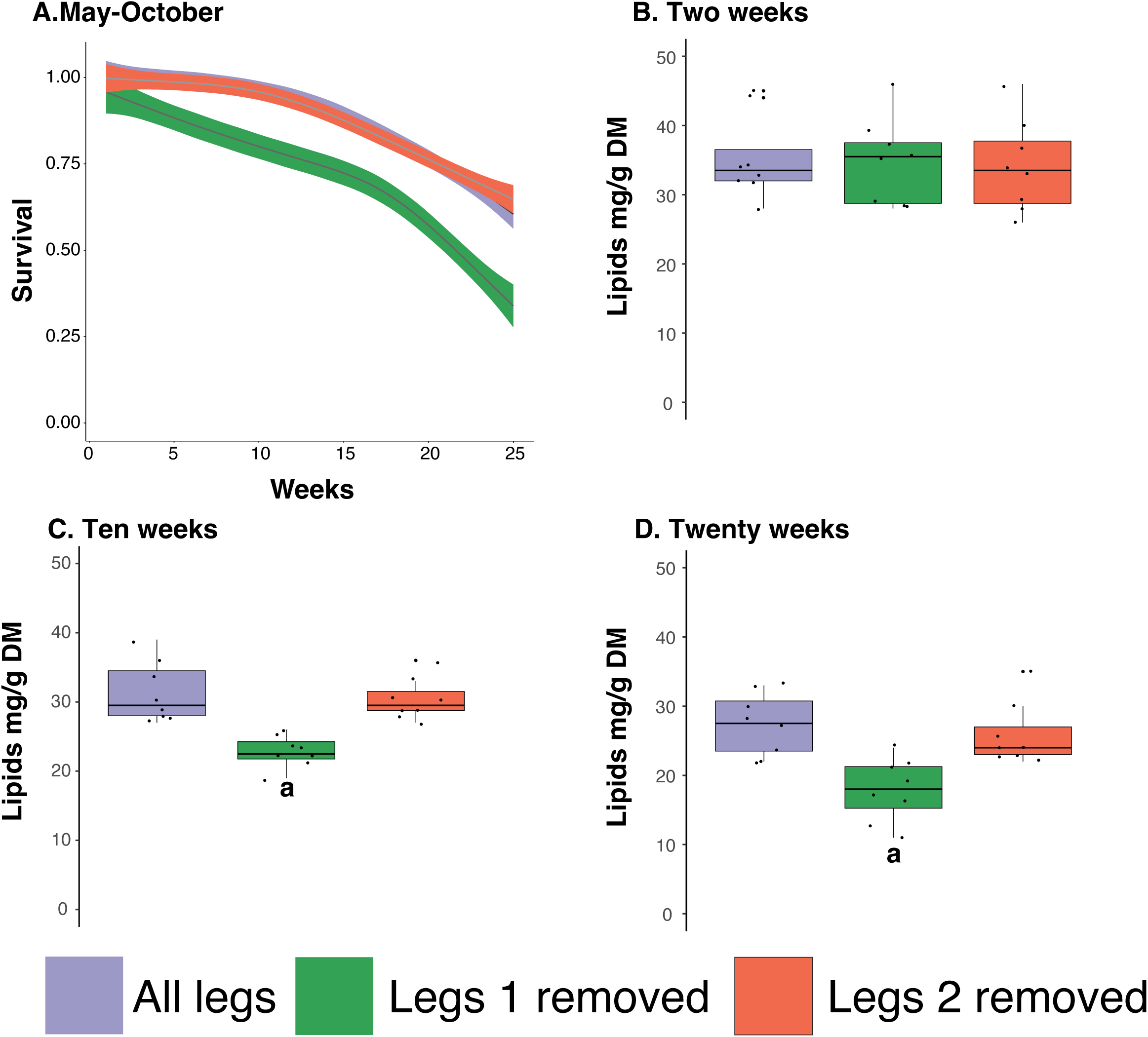
Survival and lipid content of ticks under field conditions following removal of the forelegs. A. Survival analyses. Five groups of ten ticks for each treatment were placed at the University of Cincinnati Center for Field Studies (Harrison, Ohio, USA) in small cages with local organic materials. The generalized additive model for survival shows a significant between the removal of the first legs compared to the control and second leg removal (P < 0.001). B. Lipid content after two weeks. C. Lipid content after ten weeks. D. Lipid content after twenty weeks. Eight replicates of two ticks from each treatment were used for lipid assays. a, indicates a significant difference between ticks with all legs, forelegs missing, and second legs missing, P < 0.05, Tukey’s ANOVA.

### Modeling suggests the tick survival will be reduced

To determine how impaired humidity may impact the tick population, we ran the three ADTSIM scenarios. Based on our survival studies, ticks lacking the ability to detect humidity have an increased death rate of 1.24% per week. With this increased death rate for adult ticks only (assuming both males and females will have similar death rates), there was a significant decline in ticks predicted per hectare when this impaired sensing only occurred in adults (Fig. 7A). Of interest, when the same increase in mortality for larvae and nymphs was included, there was no additional increase in mortality compared to when mortality in adults alone was considered. Throughout the season, tick levels were reduced by 44%. This modeling highlights that humidity sensing will be critical for ticks to maintain their population by allowing proper hydration and delaying nutrient utilization, but the impact is likely more severe for adult stages.

**Figure 7.**
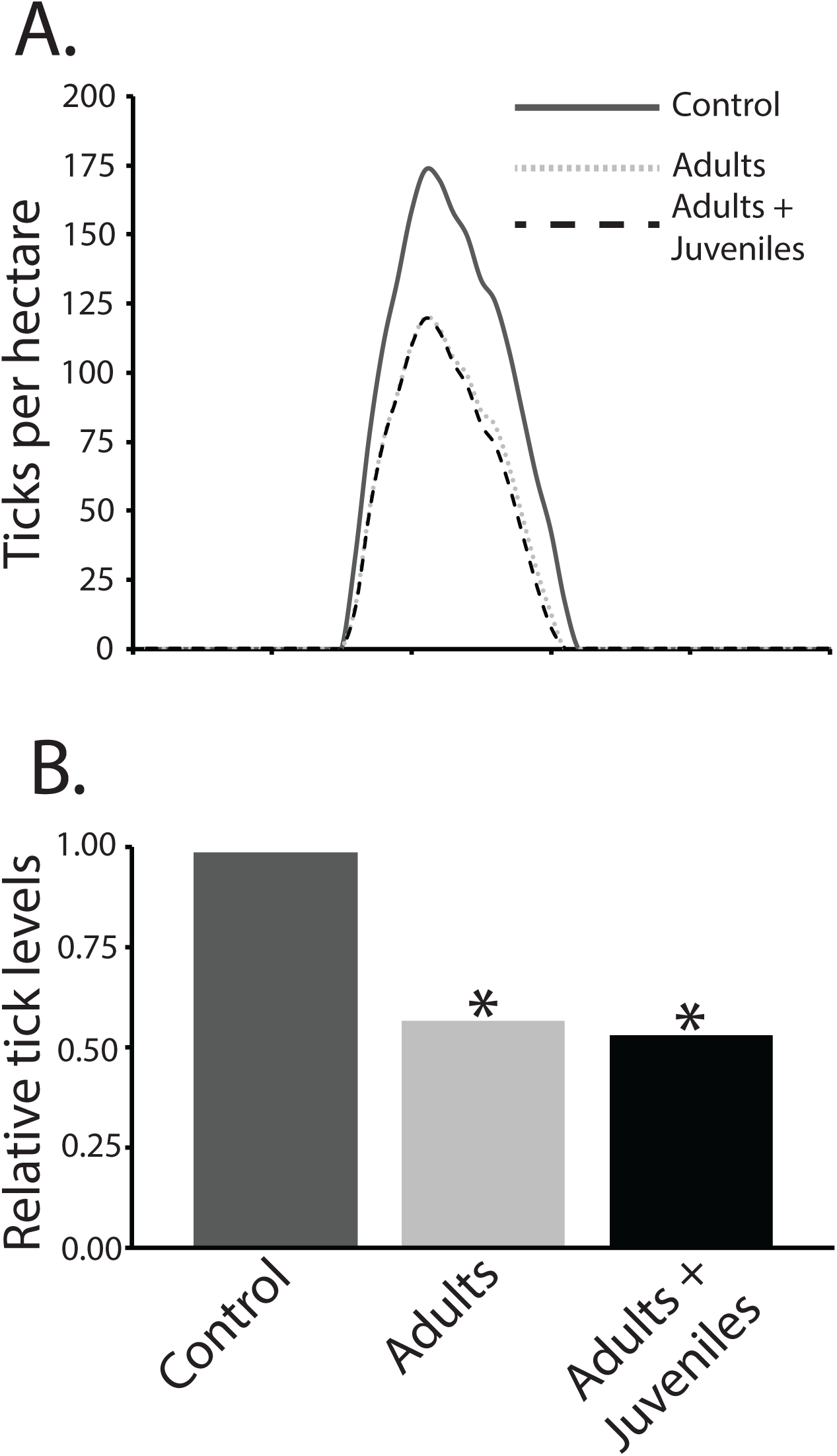
Modeling the impact of dehydration-induced mortality on tick populations. A. Ticks per hectare. Population levels were determined according to a previous model for the American dog tick (Mount and Haile, 1989) and modified according to new methods developed for *Ixodes scapularis* (Gaff et al., 2020; Sanderson et al., 2024). Environmental conditions were estimated based on those in Norfolk, VA, USA. B. Relative weekly tick levels across weeks where ticks were present. *, indicated significantly different compared to control. No differences were noted between increased mortality in adults or adults + juvenile stages. Adults and adults + juveniles were modeled with an increased mortality of 1.24% per week based on laboratory and field trial data collected in this study.

## Discussion

These studies reveal that humidity detection in ticks likely occurs in their forelegs, specifically through the Haller’s organ. This was expected as previous studies have denoted increased expression of Ir93a, a critical receptor in humidity detection (Laursen et al., 2023), in this appendage (Josek et al., 2018). Following foreleg interference, there was an increase in mortality that extended for weeks to months, particularly when allowed the choice of dry or wet conditions. This increased mortality is likely the result of chronic dehydration periods, leading to dehydration-induced mortality and increased utilization of nutrient reserves through dehydration-hydration bouts. Field observations suggest a similar mechanism, where mortality was increased over months, and ticks showed a decline in lipid levels. Lastly, modeling increased mortality yielded a substantial decline in tick populations over time. These studies indicate that humidity detection is critical for prolonged survival during off-host periods, both in remaining hydrated and preventing the elevated utilization of nutrient reserves.

Ticks must locate specific relative humidities to remain hydrated (Needham and Teel, 1991; Yoder et al., 2012). When ticks are below a specific relative humidity, dehydration will occur, which is a significant detriment to their survival (Benoit and Oyen, 2021; Rosendale et al., 2017). Most ticks can tolerate a loss of 25-35% of their water content before succumbing to dehydration stress (Benoit et al., 2007; Rosendale et al., 2016; Yoder et al., 2012). After dehydration, individuals can hydrate if a tick can locate a relative humidity above its CEH (usually above 93% RH for most tick adults) (Gaede and Knülle, 1987; Rosendale et al., 2016). Importantly, uptake is an active process, where each bout of dehydration/hydration is energetically costly (Fielden and Lighton, 1996; Rosendale et al., 2017). Both increased O_2_ consumption and CO_2_ release have been noted in relation to dehydration and hydration bouts (Fielden and Lighton, 1996; Rosendale et al., 2017). Of interest is that dehydration stress leads to metabolic expenditure that directly correlates to the severity of dehydration. Furthermore, cycling between dry and wet conditions increases metabolic rate beyond that of only dehydration, so both the process of dehydration and hydration are energetically costly. This suggests that high water loss rates and subsequent hydration through active water uptake will have a metabolic toll that will eventually lead to tick death (Fielden and Lighton, 1996; Rosendale et al., 2017). As death was not immediately induced in the weeks following interference with the Haller’s organ, the decreased survival is most likely due to a combination of increased dehydration-induced mortality along with a continual increase in the breakdown of nutrient reserves as ticks cannot locate conditions above the CEH.

Locating water for hydration or to prevent dehydration in humid microhabitats is critical to the survival of small terrestrial arthropods (Benoit et al., 2023; Hadley, 1994). Humidity detection has been characterized in fruit fly and mosquito systems (Enjin et al., 2016; Knecht et al., 2017; Laursen et al., 2023; Liu et al., 2007), where specific IRs are involved in humidity detection. Recent studies have identified that a significant contributing factor for humidity detection is IR93a, which was shown to be present in multiple tick species (Laursen et al., 2023). Here, we show that Ir93a is highly expressed in the forelegs and expression is increased during dehydration, indicating the likely enrichment in the Haller’s organ. Previous transcriptomic studies on the black-legged tick, *Ixodes scapularis*, confirm that Ir93a is highly expressed in the forelegs (Josek et al., 2018), suggesting this may be of general importance for humidity detection among all ticks. Our studies provide strong evidence that humidity sensing is accomplished through the Haller’s organ, and likely involves similar processes as identified for insect systems (Enjin et al., 2016; Laursen et al., 2023). Other specific humidity sensing receptors (Ir40a and Ir68a) have been identified in other blood feeding systems that respond to humid and dry conditions differentially (Tang et al., 2024), suggesting that humidity detection is a complex interplay between wet and dry conditions. All of these targets are well-conserved and expressed in the forelegs, suggesting a conserved role in tick humidity detection. There is evidence that sensory neurons of the Haller’s organ project to the olfactory lobe of the tick synganglion (Borges et al., 2016), but specific electrophysiology studies on ticks will be needed to confirm the specific sensilla in the Haller’s organ that is involved in humidity detection.

Outside of the inability of the ticks to sense humidity, the ticks were held under ideal conditions with little thermal stress and no predation. Even in our field trials, the ticks were placed under tree canopies, which would reduce the thermal stress as direct sunlight exposure did not occur. Under these natural conditions, which feature temperatures over 35°C for multiple days, we observe higher mortality, where 25% of deaths occurred in 25 weeks. Similar results were found in laboratory conditions after 100 weeks. The variable temperatures and humidity yield daily fluctuations in the saturation deficits that could yield dramatic increases in water loss rate (Edney, 1957; Sauer and Hair, 1971), suggesting that ecologically relevant summer conditions are likely to increase tick water loss. This is exacerbated by the inability to sense humidity, highlighted by a drastic decrease in survival under field conditions. When ticks have been observed in the field, there is significant variation in energy reserves that vary based on age (Herrmann et al., 2013), and energy stores can vary with environmental conditions experienced by each tick (Crooks and Randolph, 2006; Es et al., 1998; Randolph and Storey, 1999; Rosendale et al., 2017). These studies have indicated that physiological age, measured by energy reserves (Pool et al., 2017; Rosendale et al., 2017; Uspensky, 1995), is critical and would be directly impacted if ticks could not sense humidity, with a drastic acceleration of physiological age (Rosendale et al., 2017). Based on our estimates of lipid profiles, the lack of humidity detection would lead to a two-fold increase in physiological aging compared to those that can detect and move to more humid areas to prevent dehydration.

Following our physiological studies, modeling was used to assess how impaired humidity detection will alter population levels of ticks. The increased weekly mortality significantly impacted the population in adults, but the population was not further impacted when this increase was applied to larvae and nymphs. The lack of an additional effect in juvenile stages is likely because many other factors, such as predation and host location, are likely of more importance over the prolonged survival required for adults to have time to quest and locate a large host (Mount and Haile, 1989). Targeted studies on larvae and nymph humidity detection are likely necessary to confirm if increased weekly mortality is the same as in adults or if the juvenile stages are more sensitive to dehydration (Benoit and Oyen, 2021; Yoder et al., 2012). This may need to extend into field studies as questing patterns and other behaviors in larvae and nymphs are different than those of the adult (Benoit et al., 2021; Leal et al., 2020; Sonenshine and Roe, 2013). If weekly mortality increases by 2-3 fold, the tick population would likely decline further and impact pathogen transmission (Cooksey et al., 1990). Periods of drought and low water availability have been directly shown to reduce tick levels and, based on models, were important for tick burden (MacDonald, 2018; Mowry et al., 2019; Mowry et al., 2024; Nielebeck et al., 2023; Ogden et al., 2021). With insufficient humid detection, hydration between periods of questing in dry environments would be impaired, tick survival would likely be drastically impacted, and drought-associated reductions would yield an immediate reduction in tick populations.

### Summary

These studies suggest that impaired humidity detection will significantly impact tick populations. The importance of dry conditions during off-host periods has been established, where continual exposure to conditions below the CEH will yield a reduction in tick survival (Benoit and Denlinger, 2010; Needham and Teel, 1991). Even if ticks can survive dehydration, a bout of dehydration/hydration is energetically expensive and will deplete the finite resources as ticks do not feed between bloodmeals (McCue et al., 2017; Rosendale et al., 2017; Rosendale et al., 2019). The combined dual mortality due to dehydration and starvation would be an efficient method of reducing tick populations, where impairment of humidity detection, necessary for locating hosts and maintaining hydration, could be a target for novel control mechanisms. In addition, our studies did not target impaired humidity detection during dry periods, which could exacerbate the increased mortality we observed, as questing is likely to yield higher and more frequent water loss. Thus, if chemicals can be identified to suppress humidity detection, application during dry seasons could yield a substantial population reduction in ticks.

## Acknowledgments

Research reported in this publication was supported partially (reusable equipment) by the National Institute of Allergy and Infectious Diseases under Award Numbers R01AI148551, R21AI166633, and R21AI176098 (JBB). This work was also supported by USDA APHIS Agmt. AP22VSSP0000C033 (HDG, SS). Mention of trade names or commercial products in this publication is solely to provide specific information and does not imply recommendation or endorsement by the USDA. The USDA is an equal opportunity provider and employer.

## References

1. Bailey, S. T., Kondragunta, A., Choi, H. A., Han, J., Rotenberg, D., Ullman, D. E. and Benoit, J. B. (2023). Dehydration yields distinct transcriptional shifts associated with glycogen metabolism and increases feeding in the western flower thrips, *Frankliniella occidentalis*. Entomol. Exp. Appl. 172, 154–167.

2. Benoit, J. B. and Denlinger, D. L. (2010). Meeting the challenges of on-host and off-host water balance in blood-feeding arthropods. J. Insect Physiol. 56, 1366–1376.

3. Benoit, J. B. and Oyen, K. J. (2021). Drought and tick dynamics during climate change. In Climate, Ticks and Disease (ed. Nuttall, P.), .

4. Benoit, J. B., Yoder, J. A., Lopez-Martinez, G., Elnitsky, M. A., Lee, R. E., Jr and Denlinger, D. L. (2007). Habitat requirements of the seabird tick, *Ixodes uriae* (Acari: Ixodidae), from the Antarctic Peninsula in relation to water balance characteristics of eggs, nonfed and engorged stages. J. Comp. Physiol. B 177, 205–215.

5. Benoit, J. B., Oyen, K., Finch, G., Gantz, J. D., Wendeln, K., Arya, T. and Lee, R. E., Jr (2021). Cold hardening improves larval tick questing under low temperatures at the expense of longevity. Comp. Biochem. Physiol. A Mol. Integr. Physiol. 257, 110966.

6. Benoit, J. B., McCluney, K. E., DeGennaro, M. J. and Dow, J. A. T. (2023). Dehydration dynamics in terrestrial arthropods: from water sensing to trophic interactions. Annu. Rev. Entomol. 68, 129–149.

7. Borges, L. M. F., Li, A. Y., Olafson, P. U., Renthal, R., Bauchan, G. R., Lohmeyer, K. H. and León, A. A. P. de (2016). Neuronal projections from the Haller’s organ and palp sensilla to the synganglion of Amblyomma americanum§. Rev. Bras. Parasitol. Vet. 0, 0.

8. Carr, A. L., Mitchell, R. D., III, Dhammi, A., Bissinger, B. W., Sonenshine, D. E. and Roe, R. M. (2017). Tick Haller’s organ, a new paradigm for arthropod olfaction: how ticks differ from insects. Int. J. Mol. Sci. 18,.

9. Cooksey, L. M., Haile, D. G. and Mount, G. A. (1990). Computer simulation of Rocky Mountain spotted fever transmission by the American dog tick (Acari: Ixodidae). J. Med. Entomol. 27, 671–680.

10. Crooks, E. and Randolph, S. E. (2006). Walking by *Ixodes ricinus* ticks: intrinsic and extrinsic factors determine the attraction of moisture or host odour. J. Exp. Biol. 209, 2138–2142.

11. Edney, E. B. (1957). The Water Relations of Terrestrial Arthropods. The University Press.

12. Enjin, A., Zaharieva, E. E., Frank, D. D., Mansourian, S., Suh, G. S. B., Gallio, M. and Stensmyr, M. C. (2016). Humidity sensing in *Drosophila*. Curr. Biol. 26, 1352–1358.

13. Es, R., Hillerton, J. and Gettinby, G. (1998). Lipid consumption in Ixodes ricinus (Acari: Ixodidae): temperature and potential longevity. Bull. Entomol. Res. 88, 567–573.

14. Fielden, L. J. and Lighton, J. R. B. (1996). Effects of water stress and relative humidity on ventilation in the tick *Dermacentor andersoni* (Acari: Ixodidae). Physiol. Zool. 69, 599–617.

15. Finch, G., Nandyal, S., Perretta, C., Davies, B., Rosendale, A. J., Holmes, C. J., Gantz, J. D., Spacht, D. E., Bailey, S. T., Chen, X., et al. (2020). Multi-level analysis of reproduction in an Antarctic midge identifies female and male accessory gland products that are altered by larval stress and impact progeny viability. Sci. Rep. 10, 19791.

16. Gaede, K. and Knülle, W. (1987). Water vapour uptake from the atmosphere and critical equilibrium humidity of a feather mite. Exp. Appl. Acarol. 3, 45–52.

17. Gaff, H., Eisen, R. J., Eisen, L., Nadolny, R., Bjork, J. and Monaghan, A. J. (2020). LYMESIM 2.0: an updated simulation of blacklegged tick (Acari: Ixodidae) population dynamics and enzootic transmission of *Borrelia burgdorferi* (Spirochaetales: Spirochaetaceae). J. Med. Entomol. 57, 715–727.

18. Graeve, A., Huster, J., Mayweg, J., Fiedler, R., Plaßmann, J., Wahle, P. and Weiss, L. C. (2022). Functional assessment of the ionotropic receptors IR25a and IR93a in predator perception in the freshwater crustacean *Daphnia* using RNA interference. bioRxiv 2022.02.11.480022.

19. Hadley, N. F. (1994). Water Relations of Terrestrial Arthropods. CUP Archive.

20. Herrmann, C., Gern, L. and Voordouw, M. J. (2013). Species co-occurrence patterns among Lyme borreliosis pathogens in the tick vector Ixodes ricinus. Appl. Environ. Microbiol. 79, 7273–7280.

21. Josek, T., Walden, K. K. O., Allan, B. F., Alleyne, M. and Robertson, H. M. (2018). A foreleg transcriptome for *Ixodes scapularis* ticks: Candidates for chemoreceptors and binding proteins that might be expressed in the sensory Haller’s organ. Ticks Tick Borne Dis. 9, 1317–1327.

22. Josek, T., Sperrazza, J., Alleyne, M. and Syed, Z. (2021). Neurophysiological and behavioral responses of blacklegged ticks to host odors. J. Insect Physiol. 128, 104175.

23. Knecht, Z. A., Silbering, A. F., Cruz, J., Yang, L., Croset, V., Benton, R. and Garrity, P. A. (2017). Ionotropic Receptor-dependent moist and dry cells control hygrosensation in *Drosophila*. Elife 6,.

24. Laursen, W. J., Budelli, G., Tang, R., Chang, E. C., Busby, R., Shankar, S., Gerber, R., Greppi, C., Albuquerque, R. and Garrity, P. A. (2023). Humidity sensors that alert mosquitoes to nearby hosts and egg-laying sites. Neuron.

25. Leal, B., Zamora, E., Fuentes, A., Thomas, D. B. and Dearth, R. K. (2020). Questing by tick larvae (Acari: Ixodidae): a review of the influences that affect off-host survival. Ann. Entomol. Soc. Am. 113, 425–438.

26. Lenth, R. (2020). R. Lenth, emmeans: estimated marginal means, aka least-squares means. R package version 1.4. 5.

27. Lighton, J. R. B. and Fielden, L. J. (1995). Mass scaling of standard metabolism in ticks: a valid case of low metabolic rates in sit-and-wait strategists. Physiol. Zool. 68, 43–62.

28. Liu, L., Li, Y., Wang, R., Yin, C., Dong, Q., Hing, H., Kim, C. and Welsh, M. J. (2007). *Drosophila* hygrosensation requires the TRP channels water witch and nanchung. Nature 450, 294–298.

29. MacDonald, A. J. (2018). Abiotic and habitat drivers of tick vector abundance, diversity, phenology and human encounter risk in southern California. PLoS One 13, e0201665.

30. Maldonado-Ruiz, L. P., Park, Y. and Zurek, L. (2020). Liquid water intake of the lone star tick, *Amblyomma americanum*: Implications for tick survival and management. Sci. Rep. 10, 6000.

31. McCue, M. D., Terblanche, J. S. and Benoit, J. B. (2017). Learning to starve: impacts of food limitation beyond the stress period. J. Exp. Biol. 220, 4330–4338.

32. Mount, G. A. and Haile, D. G. (1989). Computer simulation of population dynamics of the American dog tick (Acari: Ixodidae). J. Med. Entomol. 26, 60–76.

33. Mowry, S., Keesing, F., Fischhoff, I. R. and Ostfeld, R. S. (2019). Predicting larval tick burden on white-footed mice with an artificial neural network. Ecol. Inform. 52, 150–158.

34. Mowry, S., Moore, S., Achee, N. L., Fustec, B. and Perkins, T. A. (2024). Improving distribution models of sparsely documented disease vectors by incorporating information on related species via joint modeling. Ecography.

35. Needham, G. R. and Teel, P. D. (1991). Off-host physiological ecology of ixodid ticks. Annu. Rev. Entomol. 36, 659–681.

36. Nielebeck, C., Kim, S. H., Pepe, A., Himes, L., Miller, Z., Zummo, S., Tang, M. and Monzón, J. D. (2023). Climatic stress decreases tick survival but increases rate of host-seeking behavior. Ecosphere 14,.

37. Nieto, N. C., Holmes, E. A. and Foley, J. E. (2010). Survival rates of immature *Ixodes pacificus* (Acari: Ixodidae) ticks estimated using field-placed enclosures. J. Vector Ecol. 35, 43–49.

38. Ogden, N. H., Ben Beard, C., Ginsberg, H. S. and Tsao, J. I. (2021). Possible effects of climate change on ixodid ticks and the pathogens they transmit: predictions and observations. J. Med. Entomol. 58, 1536–1545.

39. Pool, J. R., Petronglo, J. R., Falco, R. C. and Daniels, T. J. (2017). Energy usage of known-age blacklegged ticks (Acari: Ixodidae): what is the best method for determining physiological age? J. Med. Entomol. 54, 949–956.

40. Randolph, S. E. and Storey, K. (1999). Impact of microclimate on immature tick-rodent host interactions (Acari: Ixodidae): implications for parasite transmission. J. Med. Entomol. 36, 741–748.

41. Rosendale, A. J., Romick-Rosendale, L. E., Watanabe, M., Dunlevy, M. E. and Benoit, J. B. (2016). Mechanistic underpinnings of dehydration stress in the American dog tick revealed through RNA-Seq and metabolomics. J. Exp. Biol. 219, 1808–1819.

42. Rosendale, A. J., Dunlevy, M. E., Fieler, A. M., Farrow, D. W., Davies, B. and Benoit, J. B. (2017). Dehydration and starvation yield energetic consequences that affect survival of the American dog tick. J. Insect Physiol. 101, 39–46.

43. Rosendale, A. J., Dunlevy, M. E., McCue, M. D. and Benoit, J. B. (2019). Progressive behavioural, physiological and transcriptomic shifts over the course of prolonged starvation in ticks. Mol. Ecol. 28, 49–65.

44. Rosendale, A. J., Leonard, R. K., Patterson, I. W., Arya, T., Uhran, M. R. and Benoit, J. B. (2022). Metabolomic and transcriptomic responses of ticks during recovery from cold shock reveal mechanisms of survival. J. Exp. Biol. 225,.

45. Sanderson, S., Dubovyk, O., Ryan, S. J., C., W., M., Z. and Gaff, H. (2024). ADTSIM 2.0: An updated simulation of American dog tick (*Dermacentor variabilis* Acari:Ixodidae) population dynamics. In Preparation.

46. Sauer, J. R. and Hair, J. A. (1971). Water balance in the lone star tick (Acarina: Ixodidae): the effects of relative humidity and temperature on weight changes and total water content. J. Med. Entomol. 8, 479–485.

47. Sonenshine, D. E. and Roe, R. M. (2013). Ticks, people, and animals. Biology of Ticks Volume 1 1.

48. Tang, R., Busby, R., Laursen, W. J., T Keane, G. and Garrity, P. A. (2024). Functional dissection of mosquito humidity sensing reveals distinct Dry and Moist Cell contributions to blood feeding and oviposition. Proc. Natl. Acad. Sci. U. S. A. 121, e2407394121.

49. Uspensky, I. (1995). Physiological age of ixodid ticks: aspects of its determination and application. J. Med. Entomol. 32, 751–764.

50. Van Handel, E. (1985). Rapid determination of total lipids in mosquitoes. J. Am. Mosq. Control Assoc. 1, 302–304.

51. Winston, P. W. and Bates, D. H. (1960). Saturated Solutions For the Control of Humidity in Biological Research. Ecology 41, 232–237.

52. Wood, S. N. (2004). Stable and Efficient Multiple Smoothing Parameter Estimation for Generalized Additive Models. J. Am. Stat. Assoc. 99, 673–686.

53. Wood, S. (2006). Generalized Additive Models: An Introduction with R. CRC Press.

54. Wood, S. N. (2017). Generalized Additive Models: An Introduction with R, Second Edition. CRC Press.

55. Yoder, J. A., Hedges, B. Z. and Benoit, J. B. (2012). Water balance of the American dog tick, *Dermacentor variabilis*, throughout its development with comparative observations between field-collected and laboratory-reared ticks. Int. J. Acarology 38, 334–343.

