## Supplementary figures and images for "Impaired humidity sensing reduces tick survival by preventing water homeostasis"

### Supplemental Figure 1

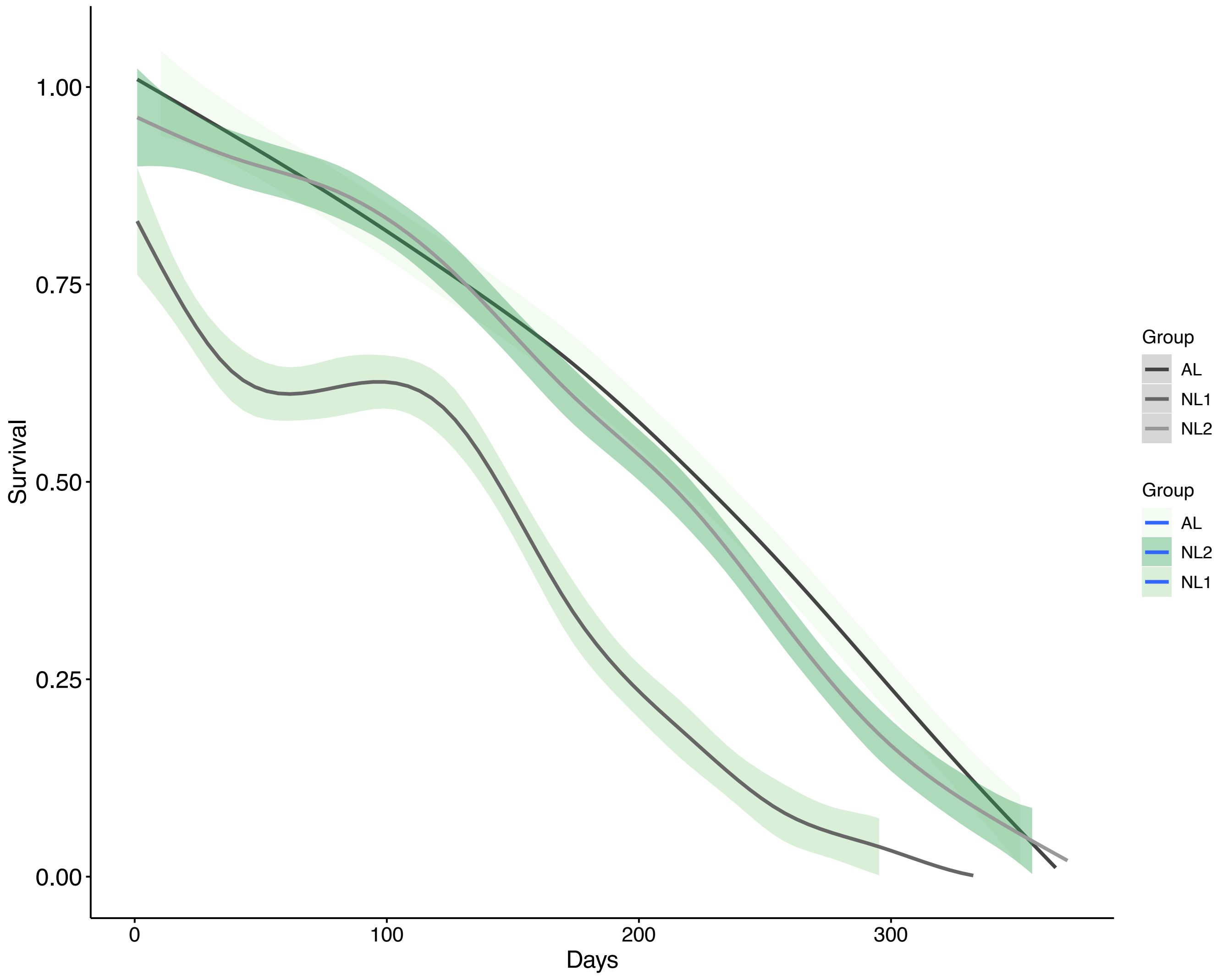

### Supplemental Figure 2

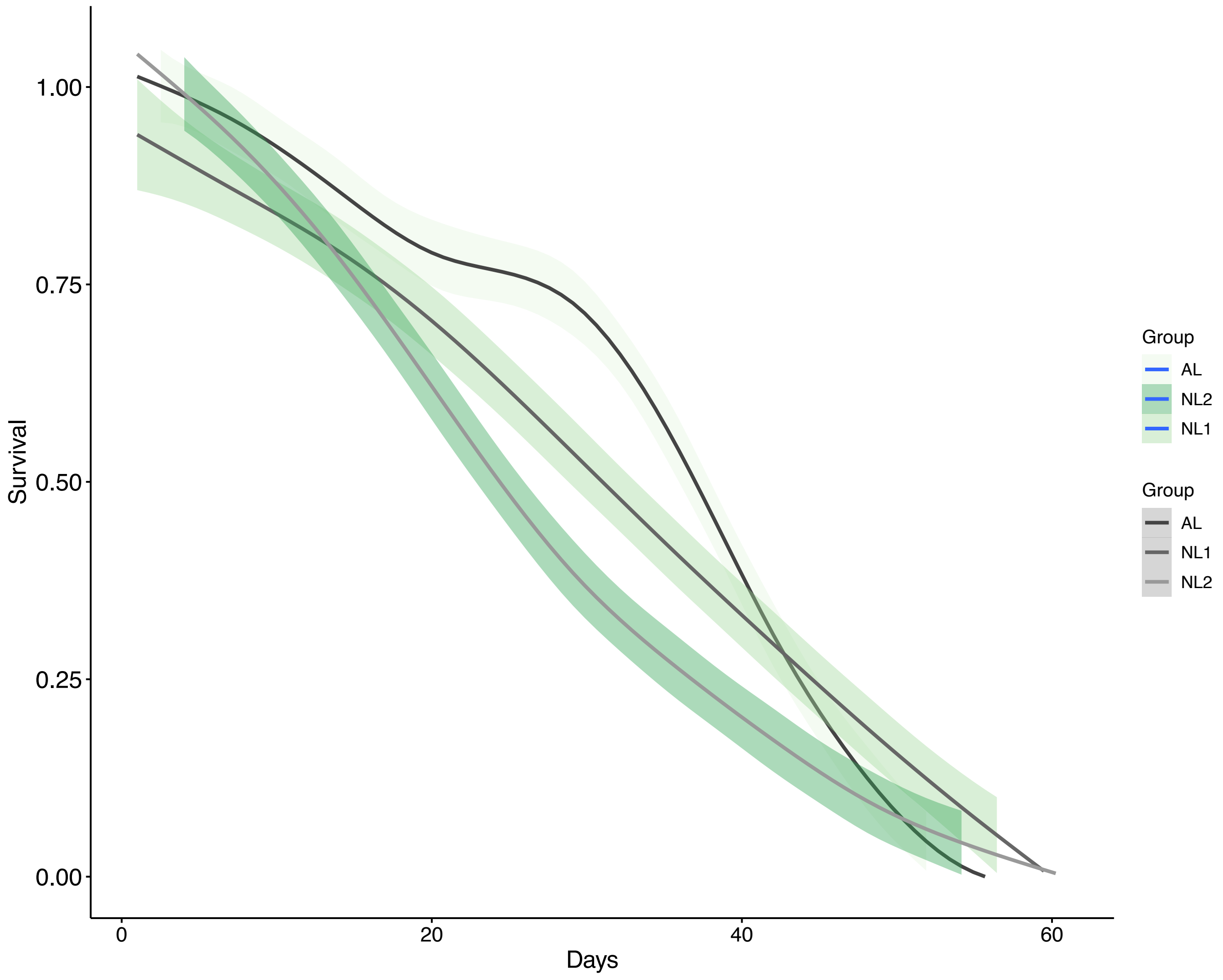
